# Human IAPP is a contributor to painful diabetic peripheral neuropathy

**DOI:** 10.1101/2021.12.03.471098

**Authors:** Mohammed M. H. Albariqi, Sabine Versteeg, Elisabeth M. Brakkee, J. Henk Coert, Barend O. W. Elenbaas, Judith Prado, C. Erik Hack, Jo W. M. Höppener, Niels Eijkelkamp

**Author notes:** **<u>Corresponding Author</u>** Dr. N. Eijkelkamp, Associate professor, Lundlaan 6, 3584 EA Utrecht, The Netherlands. Former name: Mohammed M. H. Asiri.

## Abstract

Peripheral neuropathy is a frequent complication of type 2 diabetes mellitus (T2DM). We investigated whether human islet amyloid polypeptide (hIAPP), which forms pathogenic aggregates that damage pancreatic islet β-cells in T2DM, is involved in T2DM-associated peripheral neuropathy. *In vitro*, hIAPP incubation with sensory neurons reduced neurite outgrowth and increased levels of mitochondrial reactive oxygen species. Transgenic hIAPP mice that have elevated plasma hIAPP levels without hyperglycemia developed peripheral neuropathy as evidenced by pain-associated behavior and reduced intra-epidermal nerve fiber (IENF) density. Similarly, hIAPP Ob/Ob mice that have hyperglycaemia in combination with elevated plasma hIAPP levels had signs of neuropathy, although more aggravated.

In wild-type mice, intraplantar and intravenous hIAPP injections induced long-lasting allodynia and decreased IENF density. Non-aggregating murine IAPP, mutated hIAPP (Pramlintide), or hIAPP with pharmacologically inhibited aggregation did not induce these effects. T2DM patients had reduced IENF density and more hIAPP oligomers in the skin compared to non-T2DM controls. Thus, we provide evidence that hIAPP aggregation is neurotoxic and mediates peripheral neuropathy in mice. The increased abundance of hIAPP aggregates in the skin of T2DM patients supports the notion that hIAPP is a potential contributor to T2DM neuropathy in humans.

**Graphic abstract:** 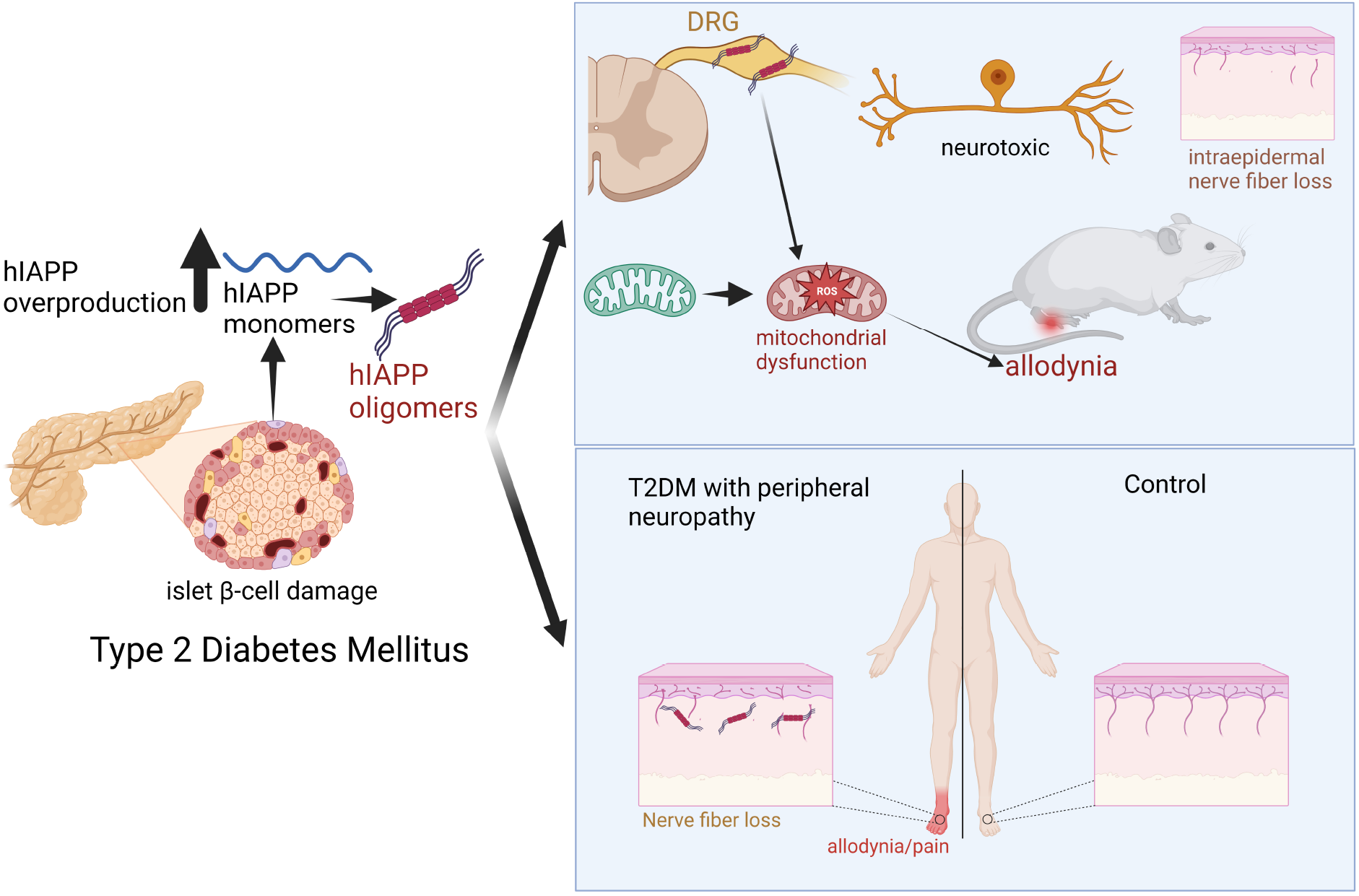

## Introduction

Diabetes mellitus (DM) is a metabolic disorder that affects around 463 million individuals globally (1). DM is one of the most significant worldwide health issues that diminishes quality of life and increases morbidity and mortality (2). Type 2 diabetes mellitus (T2DM) is the most common type of DM, characterised by insulin resistance and β-cell dysfunction (3). Diabetic peripheral neuropathy (DPN), a debilitating complication of DM, affects ∼50% of T2DM patients (4). The pathogenesis of this complication is not fully understood. Control of hyperglycaemia is not sufficient to attenuate DPN. Moreover, neuropathy can be present already in the pre-diabetic state, when hyperglycaemia has not yet developed. Thus, apparently hyperglycaemia is not the only cause of DPN (5).

Pancreatic islet amyloid is a characteristic histopathological feature of T2DM, found in ∼90% of T2DM patients. Human islet amyloid polypeptide (hIAPP) is the main component of these amyloid deposits. IAPP, or amylin, is a 37 amino acids polypeptide hormone belonging to the calcitonin gene-related peptide (CGRP) family. It is co-secreted with insulin by pancreatic islet β-cells. As a monomer, hIAPP is a soluble protein that plays a role in glucose regulation by enhancing satiety, reducing gastric emptying and suppressing glucagon release (6). Islet amyloid formation has been associated with β-cell failure in humans, monkeys, and cats with type 2 diabetes mellitus (6, 7). A hallmark of amyloid fibrils is their distinctive cross β structure, which is created by the sequential stacking of β sheets from fibril-forming protein molecules (8). Pre-fibrillar oligomer formation during protein aggregation in T2DM has been associated with cellular toxicity and a reduction in organ and cell function (9).

In T2DM, hIAPP is produced at large quantities. At high concentrations hIAPP forms toxic aggregates (oligomers and amyloid fibrils), causing β-cell death and possibly damage in other tissues (10). In contrast, IAPP from mice and rats does not form amyloid, because of a different amino acid sequence that prevents formation of toxic aggregates, and these rodents do not “spontaneously” develop T2DM (11). hIAPP amyloid deposits are not exclusively found in the pancreatic islets of T2DM patients, but are also found in other organs/tissues such as brain, heart, and kidney (12).

Considering that peripheral neuropathy is a common complication of amyloid diseases, such as familial amyloid polyneuropathy (FAP) (12), that T2DM is an amyloid disease (6, 12), and that hIAPP aggregation is not restricted to the pancreas, we investigated whether hIAPP is involved in the development of diabetic peripheral neuropathy.

## Results

To date, most research into T2DM neuropathy involves rodent models with metabolic disturbance, e.g. mice with a leptin deficiency (Ob/Ob mice). However, these models are not ideal because they lack elevated amyloidogenic IAPP and β-cell death in the pancreas as occurs in patients with T2DM when disease progresses (13).

Therefore, we evaluated whether signs of T2DM neuropathy develop in a mouse model of T2DM where hIAPP is expressed by the β-cells of pancreatic islets (hIAPP Ob/Ob). hIAPP Ob/Ob mice have significantly elevated non-fasting (Fig. 1A) and fasting (14) blood glucose levels, indicating they are diabetic. To assess development of allodynia, we used the Von Frey test, which measures the withdrawal response to innocuous mechanical stimulation. hIAPP Ob/Ob mice had allodynia from 7 weeks of age as compared to WT mice (Fig. 1B). Moreover, at 15 weeks of age, hIAPP Ob/Ob mice had reduced intra-epidermal nerve fiber (IENF) density in the plantar skin compared to WT mice (Fig. 1C/D). Thus, hIAPP Ob/Ob mice not only have metabolic characteristics of T2DM, but also show signs of diabetic neuropathy.

**Figure 1.**
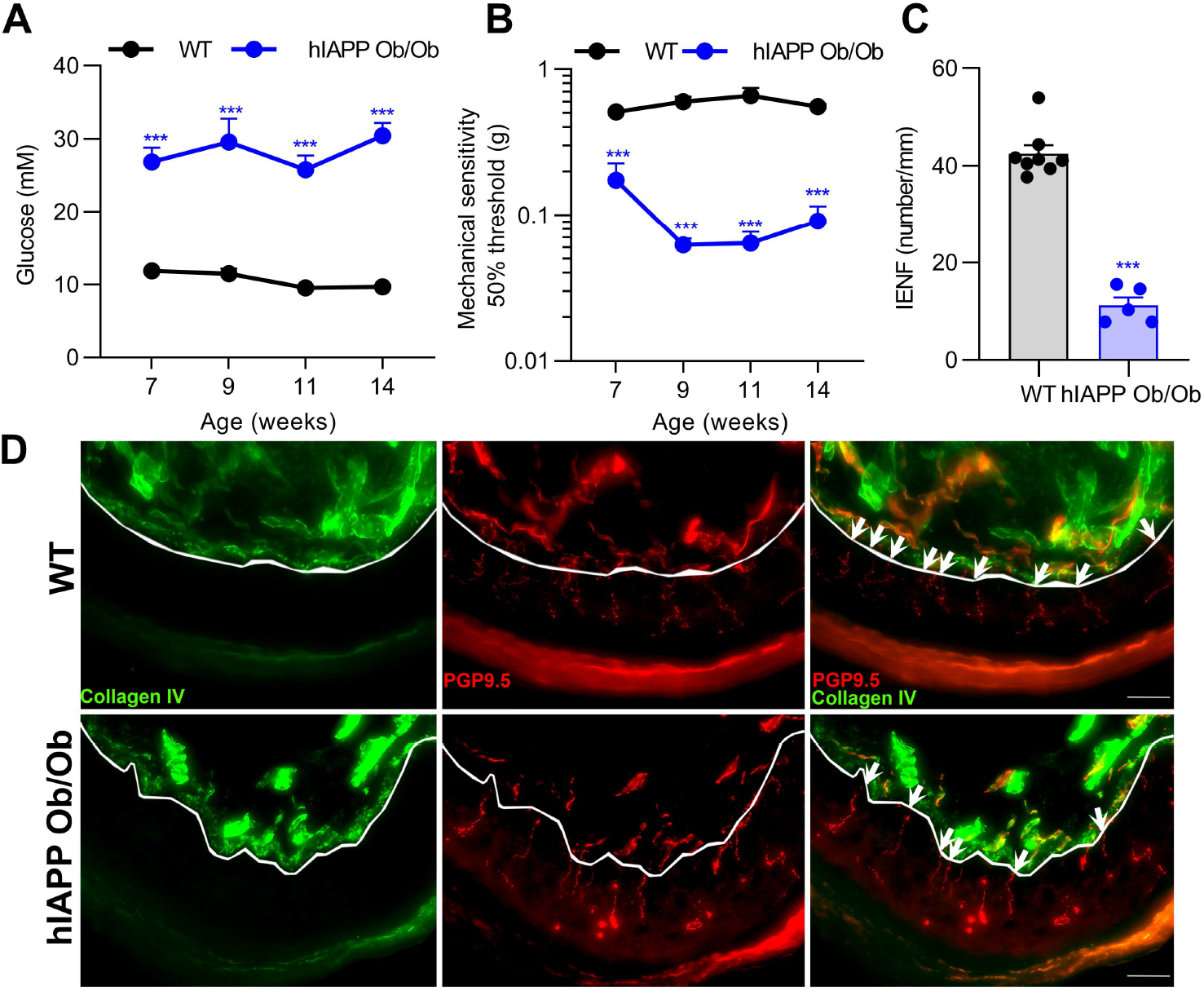
hIAPP Ob/Ob mice have features of T2DM neuropathy. (**A**) Non-fasting blood glucose levels in wild type (WT; n=8) and hIAPP Ob/Ob (n=5-8) mice; 2-way ANOVA with Sidak’s test, ***p < 0.001. (**B**) Mechanical threshold of the plantar surface of WT (n=8) and hIAPP Ob/Ob (n=7) mice; 2-way ANOVA with Sidak’s test, ***p < 0.001. (**C**) Number of nerve fibers crossing from dermis to epidermis in WT (n=8) and hIAPP Ob/Ob (n=5) mice at an age of 15 weeks; Unpaired t test, ***p < 0.001. (**D**) Representative images of paw skin of WT and hIAPP Ob/Ob mice stained for the pan neuronal marker PGP 9.5 and collagen IV (lines indicate the border between dermis and epidermis; white arrows represent intraepidermal nerve fiber; scale bar: 20 μm). All experiments were performed in male mice. Data are expressed as mean ± SEM.

Next we assessed whether hIAPP alone, thus independently of hyperglycaemia and/or obesity, is sufficient to induce signs of neuropathy. To that end, we first investigated whether hIAPP is neurotoxic by treating cultured mouse sensory neurons with hIAPP for 24 hours. Neurite outgrowth was reduced by 12% at concentrations of 100 and 1000 nM hIAPP compared to vehicle (Fig. 2A/B), without inducing cell death (Fig. S1A).

**Figure 2.**
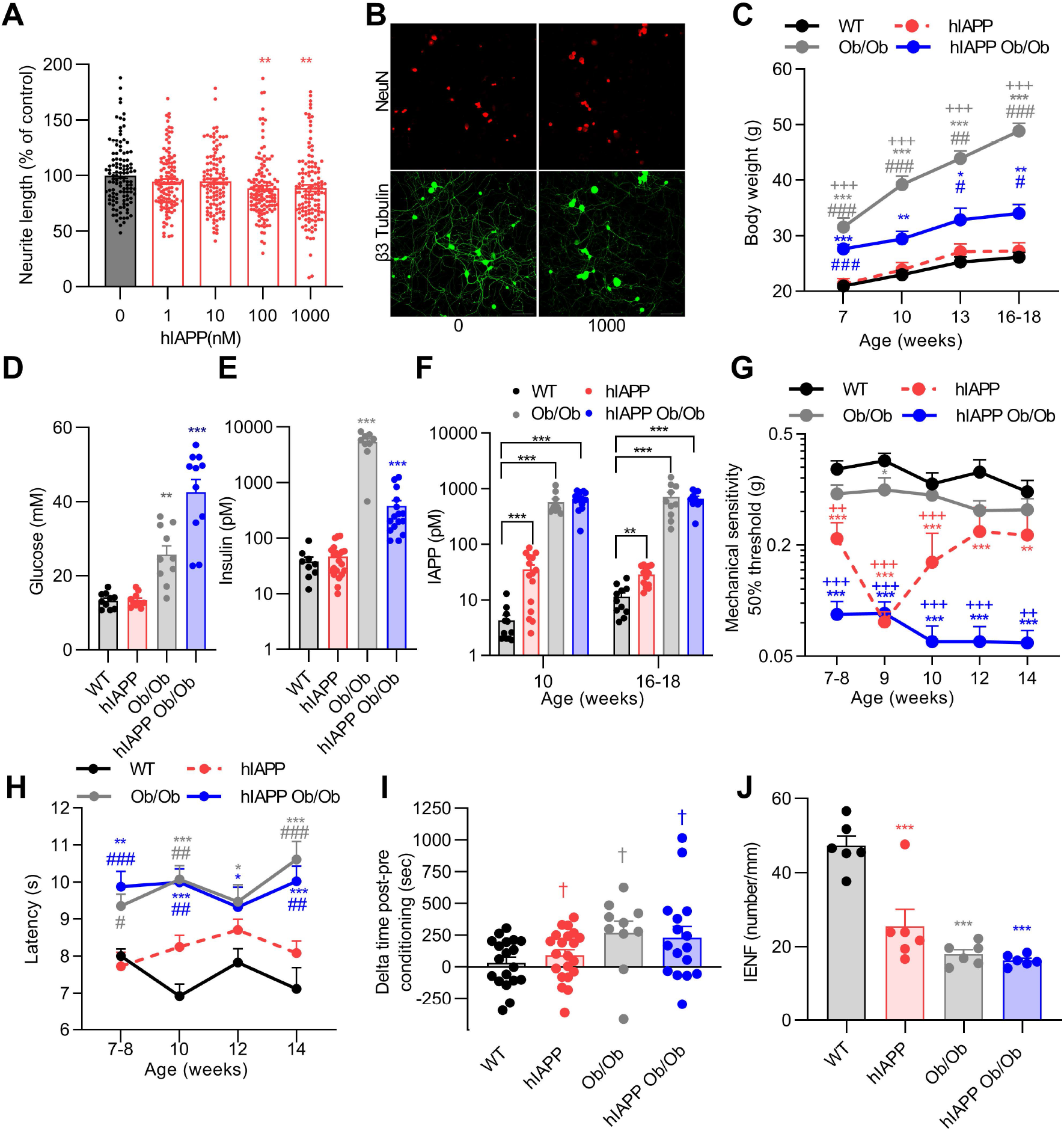
Human IAPP is neurotoxic to sensory neurons and hIAPP mice develop signs of neuropathy, also in the absence of hyperglycaemia. (**A**) Sensory neurons were treated for 24h with different concentrations of hIAPP. The average neurite length/neuron was assessed and expressed as the percentage of the average neurite length/neuron of vehicle-treated neurons (n=8; n represents a DRG culture of one mouse, male and female mice were used). (**B**) Representative images of DRG stained for β3-tubulin (green) and NeuN (red), indicating the neurite outgrowth (green), and the soma number (red); Scale bar: 50 μm. (**C**) Body weight of WT (n=12), hIAPP (n=15), Ob/Ob (n=10), and hIAPP Ob/Ob (n=15) mice. (**D**) Non-fasting plasma glucose level, (**E**) non-fasting plasma insulin level (D/E; mice age: 16-18 weeks), and (**F**) non-fasting plasma IAPP levels in WT, hIAPP, Ob/Ob and hIAPP Ob/Ob mice. (**G**) Mechanical threshold of WT (n=11), hIAPP (n=23), Ob/Ob (n=21), hIAPP Ob/Ob (n=14) mice. (**H**) Thermal sensitivity of WT (n=11), hIAPP (n=15), Ob/Ob (n= 12), hIAPP Ob/Ob (n=10) mice. (**I**) Conditioned place preference to reveal the presence of non-evoked pain. Mice were conditioned with gabapentin for 3 consecutive days and time spent in the conditioning compartment between the post and pre-conditioning phases at 15-17 weeks of age. (**J**) Quantification of IENF of the hind paw of mice at 16-18 weeks of age. All experiments were performed with male and female mice. (**A, D**,**E**) One-way ANOVA with Dunnett’s test;**p < 0.01, ***p < 0.001. (**C, F**-**H, J**) Two-way ANOVA with Tukey’s test; *p < 0.05, **p < 0.01, ***p < 0.001 vs WT; +p < 0.05, + +p < 0.01, +++p < 0.001 vs Ob/Ob mice, #p < 0.05, ##p < 0.01, ###p < 0.001 vs hIAPP (**I**) One sample t test, †p<0.05 post vs pre-conditioning. Data are expressed as mean ± SEM.

Next, we measured, over a time course of 7 to 18 weeks of age, T2DM associated parameters and pain-associated behaviors in hIAPP mice. hIAPP mice have a bodyweight, plasma glucose and plasma insulin levels comparable to those of WT mice (Fig. 2C-E). In contrast, plasma IAPP levels are increased compared to those of WT mice (Fig. 2F). The body weight of male hIAPP and WT mice was higher than in female mice (Fig. S1B). Male and female mice did not differ in other of these parameters (Fig. S1C-F). Thus, hIAPP mice do not have diabetes mellitus and provide a model to study the effects of hIAPP independently of hyperglycaemia and obesity. Intriguingly, hIAPP mice have lower mechanical thresholds as compared to WT mice (Fig. 2G). In female hIAPP mice mechanical thresholds were lower than in males (Fig. S2A). In contrast, thermal sensitivity in male and female hIAPP mice did not differ from WT mice (Fig. 2H). To further asses pain-associated behaviour, mice of ∼4 months of age were subjected to a conditioned place preference (CPP) test with the neuropathic painkiller gabapentin (15), as a measure of non-evoked pain. Male and female hIAPP mice, but not WT mice, showed place preference after conditioning with gabapentin compared to preconditioning (Fig. 2I). Overall, these data indicate that hIAPP expression is sufficient to induce pain-associated behaviours in non-obese and non-diabetic mice.

To assess whether hIAPP mice differ in their development of neuropathy compared to the more commonly used Ob/Ob mouse models, we compared hIAPP mice with obese mice (Ob/Ob), and hIAPP Ob/Ob mice. Notably, Ob/Ob and hIAPP Ob/Ob mice have increased body weight, plasma insulin levels and non-fasted plasma glucose levels (Fig. 2C-E), in line with earlier reports of elevated fasted blood glucose levels in these mice (14, 16). hIAPP Ob/Ob mice gained less body weight compared to Ob/Ob mice because hIAPP Ob/Ob mice have more severe diabetes mellitus (14) which is associated with weight loss (17). Ob/Ob mice also showed significant mechanical hypersensitivity at 9 weeks of age, but this was less severe than in hIAPP mice or hIAPP Ob/Ob mice. Whilst, hIAPP mice did not have deficits in thermal sensitivity, male and female Ob/Ob and hIAPP Ob/Ob mice had an increased latency to heat stimulation as compared to WT or hIAPP mice (Fig. 2H). Male and female mice did not differ in these parameters (Fig S2B). Ob/Ob and hIAPP Ob/Ob mice showed place preference after conditioning with gabapentin compared to preconditioning (Fig. 2I).

Peripheral neuropathy may lead to loss of IENFs. Interestingly, the number of IENFs was reduced in male and female hIAPP mice, hIAPP Ob/Ob, and Ob/Ob mice compared to WT mice (Fig. 2J). These parameters were comparable for male and female mice (Fig. S2C). Overall, these data indicate that human IAPP, in the absence of hyperglycaemia and obesity, is sufficient to induce signs of peripheral neuropathy, and this neuropathy is more severe in diabetic and obese hIAPP mice (hIAPP Ob/Ob).

Peripheral and central administration of IAPP affects the sensory nervous system in rodents and causes endothelial dysfunction, vessel wall disruption, and neurological deficits (18). To test whether hIAPP administration to WT mice is sufficient to induce pain-associated behaviors, hIAPP was injected intravenously in WT mice. A single intravenous hIAPP injection at 40 μg/kg, but not at 4 μg/kg, reduced mechanical thresholds within 2 hours of injection until 4 days after injection (Fig. 3A). The hIAPP dose that induced allodynia (40 μg/kg) results in a calculated (see methods for calculation) maximal plasma level of hIAPP that is ∼1000 times higher than in hIAPP transgenic mice (up to 100 pM) and/or humans with T2DM (up to 42 pM) (19). hIAPP has a half-life of only a few minutes (20), indicating that a short but strong rise in systemic hIAPP concentration is sufficient to cause long-lasting changes in peripheral sensory neurons. Intravenous hIAPP injection did not affect thermal sensitivity (Fig. S3A).

**Figure 3.**
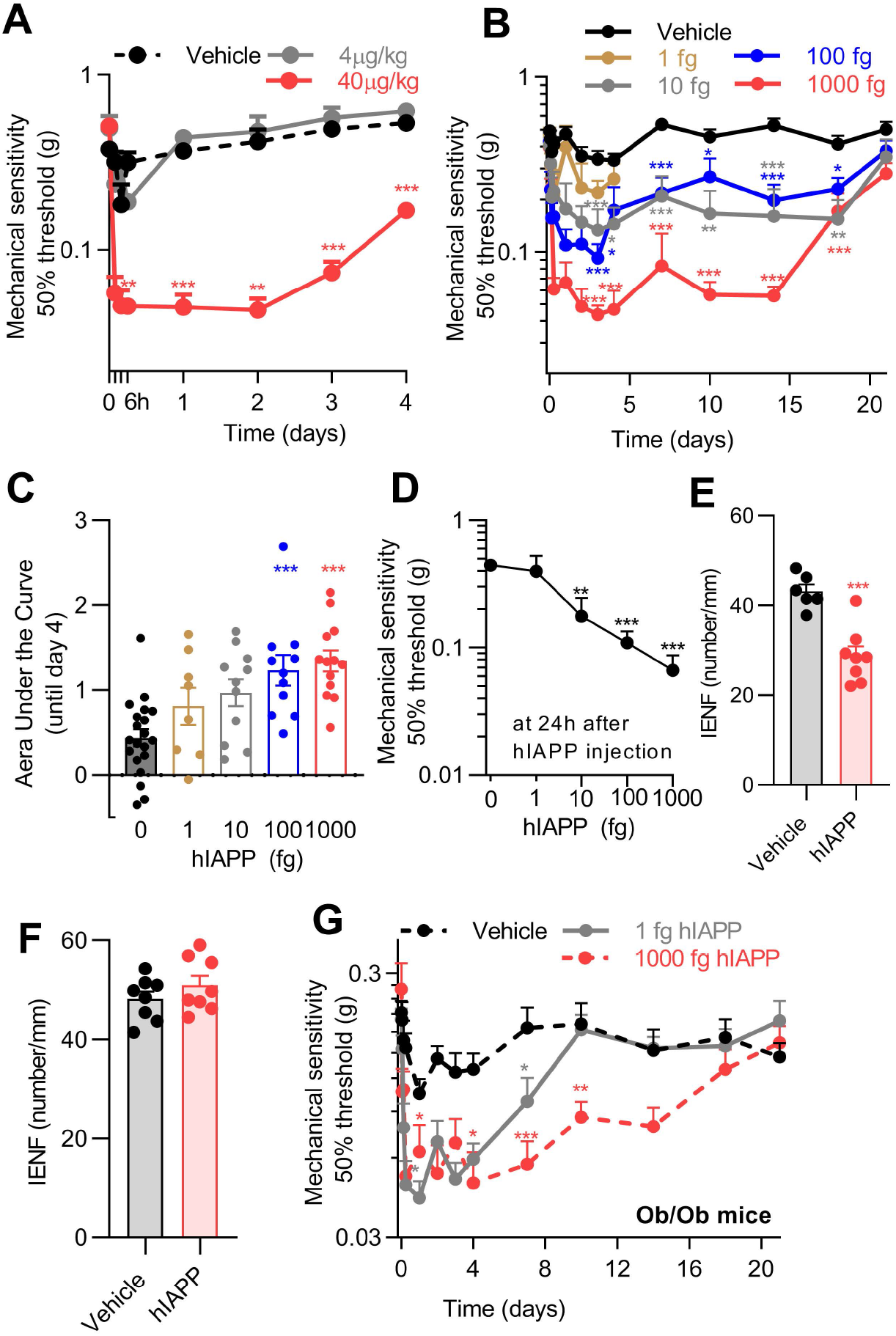
Human IAPP reduces mechanical thresholds and intraepidermal nerve fiber density in WT mice. Mechanical sensitivity of the hind paw after (**A**) intravenous (40 μg/kg, n=7; 4 μg /kg, n=3; saline, n=6) or (**B**) intraplantar injection of hIAPP (1 fg, n=8; 10 fg, n=11; 100 fg, n=11; 1000 fg, n=13; saline, n=20). (**C**) Area under the curve of the reduction in mechanical threshold from day 0-4 after intraplantar hIAPP injection of data shown in B. (**D**) Dose response curve of hIAPP-induced mechanical allodynia measured at 1 day after injection. (**E**) Quantification of plantar IENFs at 6 days after intraplantar injection of 1000 fg hIAPP or saline, or (**F**) at 27 days after hIAPP injection (1000 fg) when hypersensitivity had resolved (n=8). (**G**) Mechanical sensitivity of the hind paw after intraplantar injection of hIAPP (1 fg, n=17; 1000 fg, n=12) or saline (n=17) into male and female Ob/Ob mice. (**A**, **B, G**) Two-way ANOVA with Dunnett’s test; *p < 0.05,**p < 0.01,***p < 0.001 vs vehicle injection. (**C, D**) One-way ANOVA with Dunnett’s test; **p < 0.01, ***p < 0.001 vs vehicle injection (0 fg hIAPP). (**E, F**) Unpaired t test; ***p<0.001. Data are expressed as mean ± SEM.

Next, we tested whether local injection of hIAPP into the hind paw of WT mice induces local signs of neuropathy. Intraplantar injection of hIAPP dose-dependently induced allodynia (Fig. 3B-D). The maximum hIAPP dose (1000 fg, calculated plasma level of ∼50 pM, see methods for calculation) that elicited long lasting mechanical hypersensitivity was in the same concentration range as found in blood of hIAPP mice and T2DM patients (19). At this dose, hIAPP reduced mechanical thresholds for at least 2 weeks (Fig. 3B, S3B) and resolved within 3 weeks (Fig. S3C). IENFs measured in the plantar skin are predominantly unmyelinated (21). Intraplantar injection of 1000 fg hIAPP reduced the density of IENFs compared to vehicle injection (Fig. 3E).

Importantly, one week after hIAPP-induced hypersensitivity had resolved (Fig. S3C), the plantar skin IENF density was indistinguishable from that of vehicle-injected mice (Fig. 3F). Thus, nerve fibers recover concurrent with the resolution of hIAPP-induced mechanical hypersensitivity. Protein oligomers and larger aggregates may induce mechanical hypersensitivity through inducing inflammation. Intraplantar injection of hIAPP did not trigger expression of interleukin 6 (IL-6), IL-1β and TNF mRNA at 6 and 24 hours or 6 days after injection. hIAPP injection did increase F4/80 mRNA expression (a marker for macrophages) after one week of injection, but not at the other time points (Fig. S3D-F).

hIAPP Ob/Ob mice had stronger mechanical allodynia (Fig 2G), but also higher hIAPP levels (Fig 2F), than hIAPP or Ob/Ob mice. Thus, we questioned whether hyperglycaemic Ob/Ob mice are more sensitive to hIAPP-induced allodynia as compared to WT mice. Ob/Ob mice received 1 fg hIAPP intraplantar, a dose that did not induce allodynia in WT mice (Fig. 3B). Intriguingly, this dose increased mechanical sensitivity in Ob/Ob mice for almost 7 days (Fig. 3G). These data suggest that hyperglycaemia and obesity aggravate hIAPP-induced allodynia.

hIAPP has 43% amino acid sequence identity with hCGRP and binds to the same IAPP/CGRP receptors (22). Therefore, hIAPP may induce effects through direct actions on IAPP/CGRP receptors. In contrast to CGRP, hIAPP is toxic to rat insulinoma cells, hippocampal neurons, and astrocytes *in vitro*, independent of receptors (23, 24). To evaluate whether hIAPP-induced allodynia requires IAPP/CGRP receptors, we blocked these receptors by systemic and local administration of the IAPP/CGRP receptor antagonist CGRP 8-37, (Fig. 4A). CGRP 8-37 completely blocked the development of CGRP-induced allodynia. In contrast, the same dosing schedule of CGRP 8-37 did not block hIAPP-induced allodynia (Fig. 4B).

**Figure 4.**
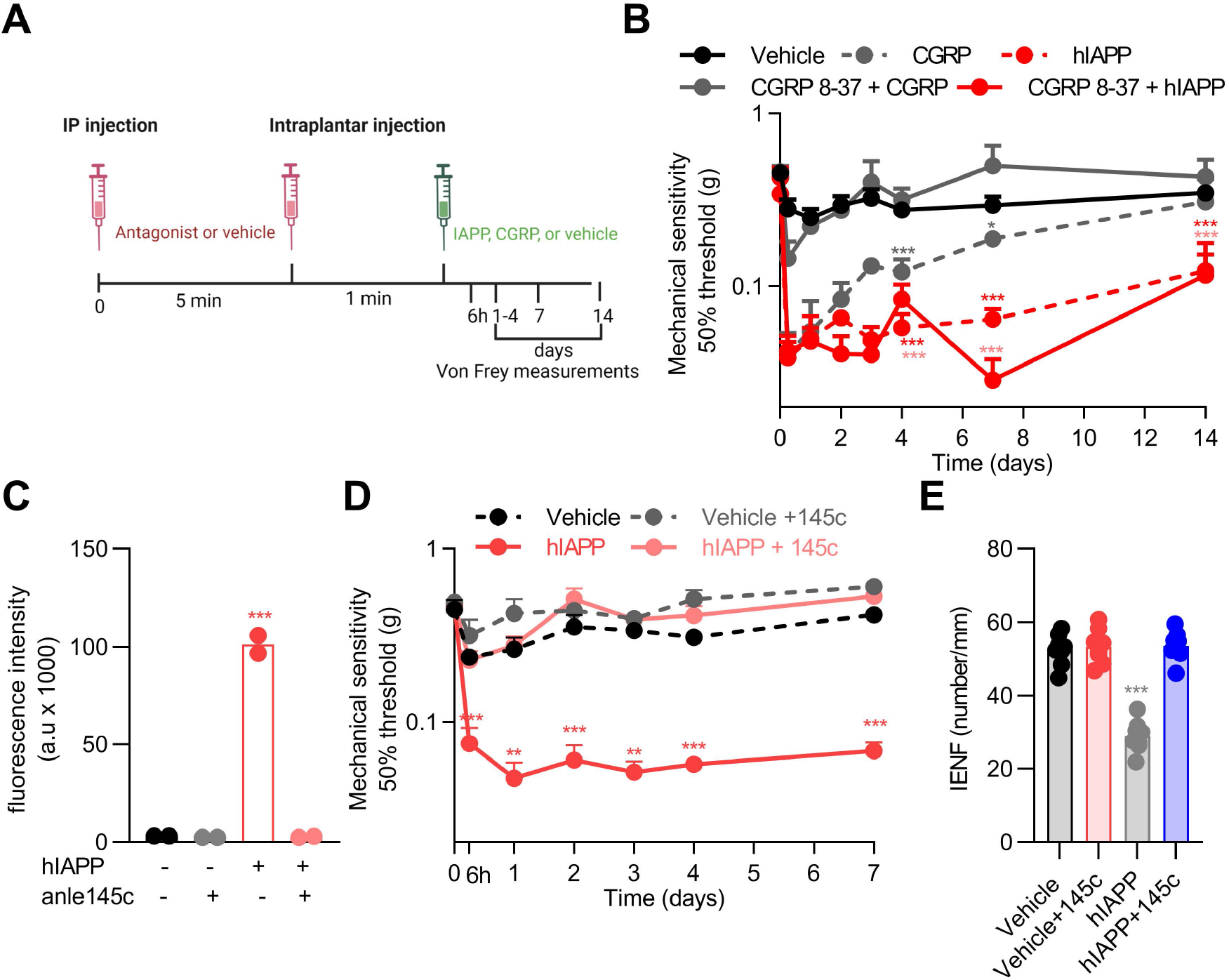
Human IAPP reduces mechanical thresholds independent of its receptors and anle 145c prevents this action in WT mice. (**A**) Schematic diagram showing the timeline and administration routes for B. (**B**) Mechanical sensitivity after antagonist (250μg/kg, CGRP 8-37) injection (n=13) or saline injection (n=13) prior to intraplantar injection of hIAPP (1000 fg, n=8) or CGRP injection (5 μg, n=5) in one paw and saline (n=13) in the other paw. Both male and female mice were used. (**C**) Fluorescence of thioflavin T after 24h. Fluorescence intensity is determined by the concentration of of amyloid fibrils. (**D**) Course of mechanical sensitivity or (**E**) plantar IENFs (n=8) at 6 days after intraplantar injection of hIAPP (1000fg/5μl) with and without anle145c (1nM), or vehicle. Treatments were prepared and incubated for 24 hrs at room temperature before injection into male and female WT mice (n=8). (**B, D, E**) Two-way ANOVA with Tukey’s test; *p < 0.05, **p < 0.01,***p < 0.001 vs vehicle. (**C**) One-way ANOVA with Dunnett’s test; *p < 0.05, ***p < 0.001 vs vehicle. Data are expressed as mean ± SEM.

To test if hIAPP aggregation is required for the development of neuropathy, we added an inhibitor of hIAPP aggregation (anle145c) (25) to hIAPP before injection. As has been observed before (25), anle145c completely inhibited hIAPP fibril formation in solution. Interestingly, anle145c completely prevented hIAPP-induced allodynia and reduction in IENF density (Fig 4C-E).

To further investigate whether aggregation of IAPP is required for its neurotoxic effects we tested mouse IAPP (non-aggregating and non-amyloidogenic) and a mutant human IAPP (Pramlintide, non-amyloidogenic) *in vitro*. Both of these non-amyloidogenic IAPP variants activate IAPP/CGRP receptor subtypes (26). Thioflavin T fluorescence assays and transmission electron microscopy showed that mIAPP indeed does not form amyloid fibrils, whereas Pramlintide only showed minor fibril formation (Fig. 5A/B). In contrast, hIAPP formed abundant amyloid fibrils (Fig. 5A/B). Treatment of cultured sensory neurons with amyloidogenic hIAPP reduced neurite outgrowth by 13% compared to vehicle control. In contrast, mIAPP or Pramlintide did not affect neurite outgrowth in sensory neurons *in vitro*. (Fig. 5C).

**Figure 5.**
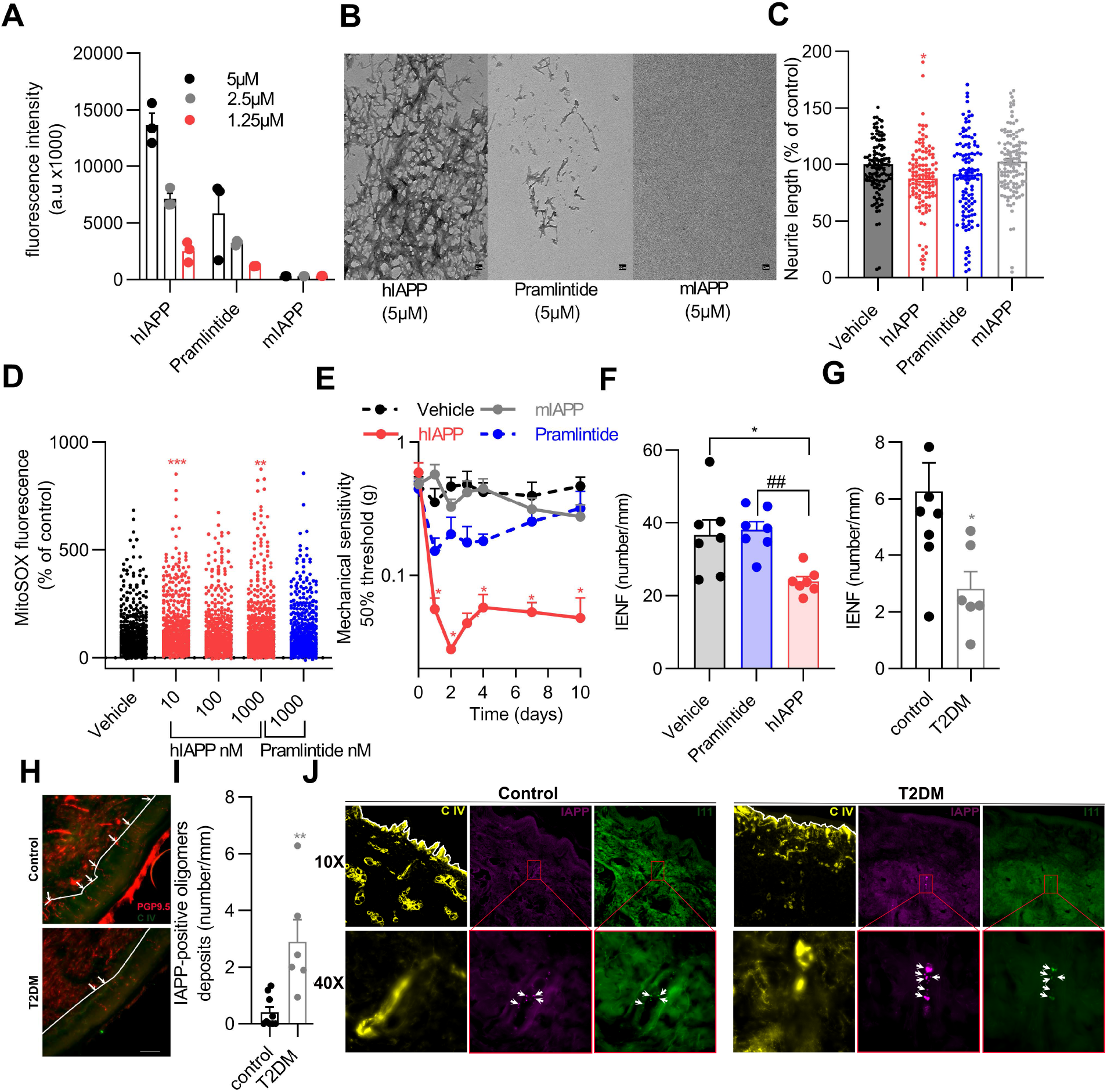
Aggregation of human IAPP is required to induce neuropathic pain. (**A**) Fluorescence (indicates amount of amyloid fibrils) of thioflavin T and (**B**) Transmission electron microscopy imaging (scale bar 0.2 nm) after 24h incubation of hIAPP, mIAPP or Pramlintide. (**C**) Sensory neurons were treated with 100 nM hIAPP, mIAPP, Pramlintide or saline for 24 hours. The average neurite length/neuron was assessed and expressed as the percentage length/neuron of vehicle-treated neurons (n=8; n represents a DRG culture of one mouse). (**D**) Mitochondrial ROS level in cultured DRG neurons incubated with hIAPP or Pramlintide (10, 100 and 1000nM). Measurements are per cell from 3 different cultures, n=551 to 654 neurons per group. (**E**) Course of mechanical sensitivity (n=5) and (**F**) IENFs density (n=7) at day 6 after intraplantar injection of 1000fg hIAPP, mIAPP and Pramlintide in male and female WT mice (**G**) Density of IENFs in skin of T2DM subjects (n=6) and none-T2DM controls (n=9). (**H**) Representative images of G stained for the pan neuronal marker PGP 9.5 and collagen IV (lines indicate the border between dermis and epidermis; white arrows represent the IENF; scale bar: 20 μm. (**I**)IAPP positive oligomers in skin of T2DM subjects (n=6) and none-T2DM controls (n=9). (**J**) Representative images of I stained for collagen IV (C IV), IAPP and oligomer staining (I11). IAPP and oligomer –positive spots are indicated by arrowheads. Lines indicate the border between dermis and epidermis. (**C, D**) One-way ANOVA with Dunnett’s test, *p < 0.05,**p < 0.01,***p < 0.001. (**E, F**) Two-way ANOVA with Tukey’s test, *P< 0.05, **p < 0.01 vs vehicle; ^##^p < 0.01 vs Pramlintide. (**G, I**) Unpaired t test; *p < 0.05, **p<0.01, ***p<0.001. Data are expressed as mean ± SEM.

Mitochondria are required for a wide range of cellular processes, including the regulation of neuronal functions (27). Mitochondria are the primary source of adenosine triphosphate (ATP) and reactive oxygen species (ROS). Mitochondria are responsible for neurotransmitter release, neuronal excitability, neuronal signaling, and plasticity. Increased ROS production in the peripheral and central nervous systems is associated with both inflammatory and neuropathic pain (27). To investigate the effect of hIAPP on mitochondrial function, cultured sensory neurons were treated with amyloidogenic hIAPP and non-amyloidogenic hIAPP (Pramlintide). Intriguingly, amyloidogenic hIAPP induced mitochondrial superoxides in sensory neurons in culture. In contrast, Pramlintide did not induce mitochondrial superoxide production in sensory neurons (Fig. 5D). Next, we determined whether IAPP aggregation is required for signs of DPN *in vivo*. Intraplantar injection of mIAPP into WT mice did not affect mechanical thresholds (Fig. 5E), whilst the same dose of hIAPP reduced mechanical thresholds for at least 10 days. Intraplantar injection of Pramlintide slightly reduced mechanical thresholds, yet this reduction was less in magnitude and duration compared to that of hIAPP (Fig. 5E). Injection of Pramlintide did not affect IENF density, whilst the same dose of hIAPP caused a reduction of ∼50% (Fig. 5F). Overall, these data indicate that the ability of IAPP to form aggregates/fibrils is required for its neurotoxicity *in vitro* (28), and to induce allodynia and reduce IENF density *in vivo*.

hIAPP can accumulate in nervous tissues, e.g. in hippocampal neurons of hIAPP transgenic mice and rats, and form aggregates that are associated with neurological deficits (29). hIAPP oligomers destabilize cell membranes and form membrane pores (30). Therefore, we assessed whether IAPP oligomers are present in the DRGs that contain the soma of sensory neurons, Interestingly, IAPP-positives oligomers were detected in DRGs of hIAPP mice but not in DRGs of WT mice (Fig S4). Next, we assessed the presence of hIAPP oligomers in the skin of T2DM patients with neuropathy (Suppl. Table l). T2DM patients had neuropathy (Suppl. Table 1) and a reduction in IENF density compared to non-T2DM controls (Fig. 5G/H). Intriguingly, T2DM patients with neuropathy had significantly more IAPP-positive oligomers in the skin compared to controls (Fig. 5I/J). These data suggest that potential toxic aggregates of hIAPP are more abundant in the skin of T2DM neuropathy patients.

## Discussion

Although the cause of neuropathy in T2DM is not fully understood, three characteristics of T2DM may contribute to the development of diabetic neuropathy, i.e. hyperglycaemia, obesity, and amyloidogenic IAPP (5, 12). Here, we present evidence that amyloidogenic hIAPP is a contributor to the development of diabetic neuropathy. Diabetic and non-diabetic mice expressing hIAPP developed signs of neuropathy such as allodynia and nerve damage in the skin. Importantly, the aggregation of IAPP is required to induce these signs of neuropathy. Thus in addition to hIAPP aggregation being associated with dysfunction and death of pancreatic β-cells in T2DM (6), hIAPP also damages peripheral sensory neurons and contributes to neuropathy development. Importantly, we found that hIAPP-positive oligomers are present in hIAPP mice and also in the skin of T2DM patients. These findings not only add to earlier findings that IAPP oligomers are found outside of the pancreas, they also support the notion that hIAPP is a contributor to diabetic neuropathy in humans.

Ob/Ob mice, with hyperglycaemia exhibit T2DM neuropathy and are often used to study T2DM neuropathy (31). Similarly other mouse models with metabolic disturbances have been used, such as the db/db mouse (32). However, these mice are not an ideal model for human T2DM because they lack human IAPP expression and the death of islet β-cells when disease progresses. hIAPP Ob/Ob mice become insulin resistant, hyperglycaemic, and develop pathogenic amyloid deposits in pancreatic islets due to overproduction of hIAPP by the islet β-cells (6, 14). Given the accumulating evidence that aggregating hIAPP contributes to development of T2DM (3), we propose that hIAPP Ob/Ob mice are a better model to study T2DM, diabetic neuropathy, and potentially other disease-associated symptoms, as compared to Ob/Ob and db/db mice.

Hyperglycaemic Ob/Ob mice were found to be more sensitive to hIAPP-induced allodynia than WT mice. Although hyperglycaemia may not be the only driver of diabetic neuropathy, it may predispose to IAPP-induced hypersensitivity because hIAPP-induced cell toxicity is exacerbated by hyperglycaemia-induced glycated insulin (33). In addition, obesity sensitizes neurons to pain-inducing stimuli (34) and hyperglycaemia, as well as obesity related insulin resistance, increases IAPP transcript levels and IAPP biosynthesis in pancreatic islet β-cells (35). Overall, these findings possibly explain why hIAPP Ob/Ob mice have higher mechanical sensitivity than non-obese hIAPP mice.

Our findings show that aggregation of hIAPP is necessary for the development of signs of neuropathy, suggesting that non-receptor mediated mechanisms are probably involved. Indeed, we identified that hIAPP -induced neuropathy does not involve the IAPP/CGRP receptor. Aggregation of IAPP is required to exert its damaging effect on neurons *in vitro* and blocking hIAPP aggregation, using the small molecule inhibitor anle145c (25), prevented hAPP-induced neuropathy *in vivo*. Pramlintide, a non-amyloidogenic hIAPP variant did show some fibril formation *in vitro* and induced some small neuropathic-like effects *in vitro* and *in vivo* in mice, but considerably less as compared to hIAPP. Since Pramlintide is used in clinical practice (36), its potential neuropathic effects warrant further investigation.

In T2DM patients, small nerve fiber pathology is observed early in the development of painful DPN, sometimes even prior to the presence of hyperglycaemia (37). In this study, we found more abundant hIAPP oligomers in the dermis of patients with T2DM neuropathy compared to non-diabetic controls. This raises the question why hIAPP depositions were found in the dermis if small nerve fibre degeneration and a reduction in IENF density are related to nerve terminal loss in the epidermis? Likely, hIAPP oligomers also harm nerve fibres in the dermis, ultimately contributing to the degeneration of nerve fibre terminals in the epidermis. In line with this rationale, different neurotoxic drugs (e.g., chemotherapeutics) target neurons at different levels, but degeneration often begins in the epidermis (38). In addition, hIAPP aggregates may exist at other areas of the peripheral sensory nervous system that we are unable to evaluate with our skin biopsies. In this context, we have demonstrated the presence of hIAPP aggregates in DRGs of hIAPP mice, but not in WT mice (Fig. S4), suggesting that similar aggregates may also arise in humans.

How hIAPP mechanistically causes neuropathy remains unclear. hIAPP aggregation, with formation of oligomers and amyloid fibrils, induces cytotoxicity of pancreatic islet β-cells through various mechanisms, including membrane disruption, impaired mitochondrial function, and autophagy malfunction (39). Many of these cellular abnormalities, especially mitochondrial defects (27, 40), have been implicated in pain conditions, the development/progression of diabetic neuropathy, as well as in other forms of amyloid neuropathy (12). We found increased mitochondrial ROS levels and reduced neurite outgrowth in hIAPP-treated sensory neurons, supporting the notion that aggregating hIAPP induces neurotoxicity through mitochondrial damage (12). These pathogenic mechanisms may contribute to DPN not only by impacting neurons, but also other cell types involved in neuropathy, such as macrophages, microglia, satellite glia cells, Schwann cells and endothelial cells (12). Future studies are needed to determine which pathogenic pathways and which cell types are involved in hIAPP-mediated DPN.

In conclusion, our data show that human IAPP induces signs of peripheral neuropathy in the absence of hyperglycaemia and aggravates mechanical allodynia in hyperglycaemic obese (Ob/Ob) mice. Therefore, inhibition of hIAPP aggregation is a novel approach to treat and/or prevent DPN, as it plays a crucial role in the progression of T2DM-associated neuropathy.

## Methods

### Animals

Power calculations were performed to estimate the sample size needed to detect a minimal predefined effect size. All animals were randomly allocated to a group before the start of any measurements or treatment. Experiments were conducted using both male and female mice aged 8 –16 weeks. Observers performing behavioral experiments were blinded with respect to treatment /genotype groups. Mice were housed in groups under a 12 h light/12 h dark regime, food and water was available ad libitum. The cages contained environmental enrichments including tissue papers and shelter opportunities. Mice were acclimatized to the experimental setup for 1-2 weeks prior to the start of each experiment. To avoid potential cage bias, animals were randomly assigned to the different groups prior to the start of experiments, treatment groups were equally divided per housing cage, and experimenters were blinded for the treatments and genotypes.

Four groups of mice were used, 1) transgenic mouse model of T2DM (Obese (Ob/Ob) mice that produce hIAPP in their pancreatic islet β-cells; hIAPP Ob/Ob), 2) non-obese hIAPP transgenic mice, 3) Ob/Ob mice, and 4) wildtype (WT) mice.

The generation of these four separate sublines from the original GG2653 line (hIAPP x Leptin Ob, on a C57Bl6 background) was previously described (14, 41). The generation of the mice was performed in two breedings: 1) Ob heterozygous (Ob/+) x Ob heterozygous (Ob/+) and 2) hIAPP homozygous/Ob heterozygous x hIAPP homozygous/Ob heterozygous.

Ob/ + mice were mated with Ob/+ mice to generate non-transgenic Ob/Ob mice (homozygotes) and WT mice. To maintain this strain, WT mice (+/+) were mated with Ob/+ mice.

hIAPP homozygous Ob/Ob mice and hIAPP homozygous non-obese mice were generated by mating hIAPP homozygous Ob/+ mice with hIAPP homozygous Ob/+ mice. The distinction between Ob/Ob, Ob/+ and WT littermates in both breedings was performed by genotyping a leptin -Ob gene PCR product of 250 base pairs (bp) using the restriction enzyme DdeI. This results in 2 DNA fragments for WT mice (150 bp and 100 bp), while the Ob point mutation creates an additional DdeI site within the 100 bp fragment, resulting in three DNA fragments after DdeI digestion for Ob/+ mice and complete loss of the 100 bp fragment for ObOb mice.

### Glucose, IAPP and insulin measurements

Blood was obtained by cheek puncture from non-fasted mice at age 10 weeks and at the end of the experiments (at an age between 15-18 weeks). Blood was collected in EDTA-tubes (Greiner, Minicollect) and saved on ice until centrifugation at 3000 rpm for 5 min at 4 °C. Plasma was taken, divided into aliquots and stored at -80 °C until analysis.

Glucose was measured in EDTA plasma using the hexokinase method.

Insulin was measured in EDTA plasma using a radio immuno assay (RIA, Merck Millipore, RI-13K). IAPP was measured by an enzyme-linked immunosorbent assay (ELISA; Merck Millipore, EZHA-52K).

### Drug dilution and administration

hIAPP, mIAPP (both obtained from Bachem, Switzerland) and Pramlintide (obtained from ANASpec, USA) were dissolved in dH2O to obtain stock solutions of 200 μg/ml; aliquots were stored at -80°C. Human calcitonin gene-related peptide α (CGRPα) and human calcitonin gene-related peptide 8–37 (CGRP 8-37) were obtained from Bachem (Switzerland) and dissolved in 0,9% NaCl (1 mg/ml) before injection. For behavioral experiments, WT male mice received intravenous injection (IV) of hIAPP (40 μg/kg, 4 μg /kg) or vehicle (0,9 % NaCl) at an age between 8-16 weeks.

The intravenous injection dose was calculated based on the estimation that WT mice have 2 ml of blood and the physiological range of plasma IAPP concentrations in healthy WT mice is 20-100 pM (28). Thus, injection of 40 μg/kg intravenously would correspond to 125nM in blood, which is ∼1000 times higher than the physiological range.

hIAPP (10 fg-1000 fg; 5 μl), mIAPP (1000 fg; 5μl, or Pramlintide (1000 fg; 5μl) was injected intraplantarly in one paw. The other paw received the same volume of 0,9 %NaCl. For intraperitoneal injections, mice received of the IAPP/CGRP antagonist (CGRP8-37, 250 μg/ kg,) or vehicle (0,9 % NaCl) at 5 minutes prior to the intraplantar injection of hIAPP. Also at 1 minute prior intraplantar injection of hIAPP (1000fg/5μl) or CGRP (5μg/5μl) mice received an intraplantar injection of CGRP8-37 (5μg/5μl).

For intraepidermal nerve fiber (IENF) quantification, WT mice received an intraplantar injection of 1000 fg hIAPP or Pramlintide in 5μl saline or vehicle. For blocking hIAPP aggregation with anle145c, monomeric hIAPP was dissolved with Hexafluoroisopropanol (HFIP) to obtain a monomeric stock solution. Anle145c was dissolved as 10mM stock solution in DMSO. A 50μM hIAPP solution was prepared from this stock in Tris 10mM, NaCl 100mM, Ph= 7.4 and incubated overnight at room temperature with or without a 20-fold molar excess of anle145c (1 mM) with 10% DMSO. This solution of hIAPP was diluted to 50 pM in saline before intraplantar administration of 5 ul to the mice

### Intraepidermal nerve fiber (IENF) quantification

Skin of hindpaw was taken from mice after euthanasia, then incubated with Zamboni fixative overnight at 4°C, rinsed overnight in 30% sucrose in PBS at 4°C and then cryoembedded in mounting media (OCT).The skin was cryosectioned at 20 μm for immunohistochemistry; sections were incubated with rabbit anti-PGP9.5 antibody (1:1000, Sigma-Aldrich, US) and goat anti-collagen IV antibody (1:200, Southern Biotech, US) for 24 hours at 4°C. Sections were then rinsed 3 times in PBS+ 0.3% triton X-100 and incubated with Alexa Fluor 594 labelled donkey anti-rabbit antibody (1:500, Life Technologies) or Alexa Fluor 488 labelled donkey anti-goat antibody (1:500, Life Technologies), respectively, for 2 hours in the dark at room temperature, and with 4,6-diamidino-2-phenylindole DAPI (1:5000) for 5 minutes, before being rinsed (2 times in distilled water) and mounted onto slides. A stack of 12 images per hind paw skin was obtained by using a Zeiss confocal microscope (×40 objective) and the number of nerves fibres that crossed from the dermis to the epidermis per linear mm of skin was quantified.

### Real-time quantitative PCR

Total RNA was isolated from hind paw skin that was isolated at day 6 after intraplantar injection of vehicle, or of 1000fg/5 µl hIAPP or Pramlintide, using Trizol (Invitrogen, Paisley, UK) and RNeasy mini kit (Qiagen, Hilden, Germany) according to manufacturer‘s protocol.

cDNA was synthesized using iScript reverse transcription supermix, in accordance with the manufacturer’s instructions (Bio-Rad, Hercules, CA). Real-time quantitative PCR was then performed with iQ SYBR Green Supermix (Invitrogen, Paisley, UK). We used an amount of cDNA corresponding to 1-5 ng of RNA input per qPCR reaction and the following primer pairs:

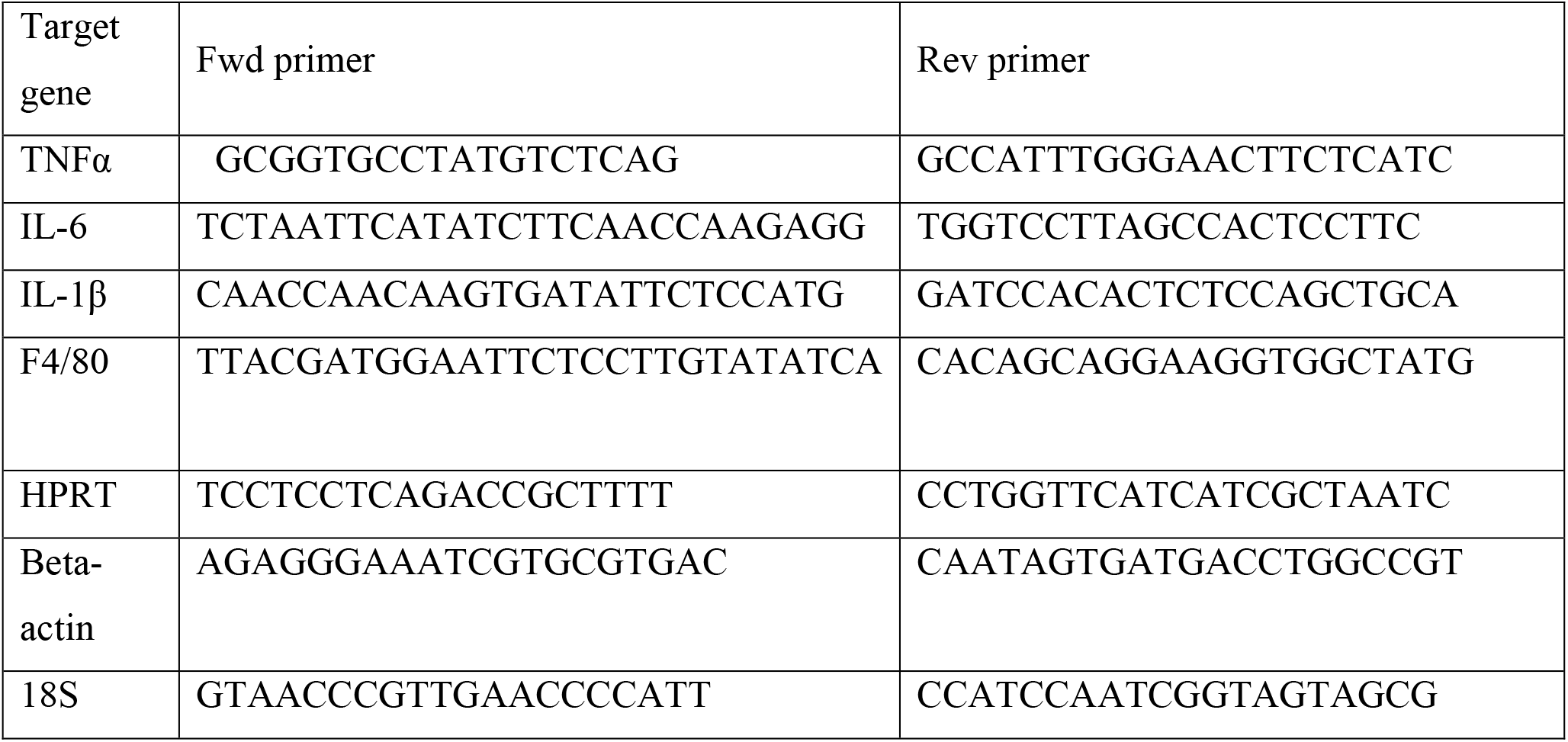

mRNA expression is represented as relative expression = 2^Ct(average of housekeeping genes (18S,HPRT, Beta actin)-Ct(target).

### Pain Measurements

Mechanical sensitivity was measured by Von Frey hairs (Stoelting, Wood Dale, IL, US), and the 50% paw-withdrawal threshold was calculated using the up-and-down method. Briefly, von Frey filaments were placed to the plantar surface of the paw for a maximum of 5 seconds. In the event of a non-response after the first filament (0.4 g), the next filament with a higher force was applied. In the event of a response, the filament with the lower force was used. A minimum of 30 seconds was taken between the application of filaments. After the first change of direction, four readings were taken (42). Thermal sensitivity was assessed by Hargreaves test (IITC Life Science). In short, the Hargreaves test was conducted in a Perspex enclosure (box) with a heated (32°C) glass bottom. Underneath the animals, a radiant heat source was focused at the plantar surface of the hind paw. The time taken until a withdrawal response from the heat stimulus was recorded as the withdrawal latency. Paws were measured at least 3 times with at least 30 seconds between measurements. In WT C57Bl/6 mice, the intensity of the light source was tuned to generate withdrawal latencies of 8-10 seconds, with a 20-second cut off to avoid tissue injury (43).

Spontaneous pain was measured by the conditioned place preference test (CPP, Stoelting, Wood Dale), as previously described (44, 45). In short, a three-chambers box (A, B, and C) is used (chamber A and C measure 18x 20 cm), with the B chamber being a smaller chamber connecting A and C divided from the neighbouring chambers by a divider with an entrance. The A and C chambers had various patterned walls and floors that were texturally different. At day 1 (pre-conditioning) animals were allowed to move freely for 30 min and the time spent in each chamber was recorded. At days 2-4, the mice were conditioned with daily intraperitoneal injections of vehicle or the pain-killer gabapentin (100 mg/kg, Sigma-Aldrich, St. Louis, MO, US). In the morning, the animal was placed for 30 min in one closed chamber (black room), 1 minutes after receiving vehicle. In the afternoon (3 hours later), the mouse was placed for 30 min in the other chamber (white room) 1 minutes after receiving gabapentin treatment. At day 5 post-conditioning the animals were placed in the start chamber (Chamber B) and allowed free access for 30 min to either chambers (A or C). The time spent in the chambers was recorded. To define treatment effect, the mean of the time spent in the treatment-associated chamber (white room) during adaptation (day 1; pre-conditioning) was subtracted from the time spent in that chamber on the test day (day 5; post-conditioning). A significantly longer stay in the white room after conditioning indicates the presence of spontaneous pain.

### Primary cell cultures

Adult mice Dorsal root ganglia (DRG) were dissected after mice were killed and then placed in ice-cold dissection medium (HBSS w/o Ca^2+^ and Mg^2+^ (Gibco 14170-088), 5mM HEPES (Gibco 15630-049), and 10mM glucose (Sigma G8769). After dissection, axons were cut off and subsequently the DRG were digested, in order to get a cell suspension, with of mixture of HBSS w/o Ca^2+^ and Mg^2+^, 5mM HEPES, 10mM glucose, 5mg/ml collagenase type XI (Sigma) and 10mg/ml Dispase (Gibco), for 30-40 minutes at 37°C in a 5% CO2 incubator. After that, Dulbecco’s modified Eagle’s medium (DMEM, Gibco 31966-021) with 10% heat-inactivated foetal bovine serum (FBS, Sigma F9665) was added to inactivate the enzyme mixture. Cells were centrifuged for 5 min at 79 g., resuspended in Dulbecco’s modified Eagle’s medium (Gibco) containing 10% FBS (Gibco), 2 mmol/L glutamine (Gibco), 10,000 IU/ ml penicillin-streptomycin (Gibco) and plated on poly-L-lysine (0.01 mg/mL; Sigma) and laminin (0.02 mg/mL; Sigma)-coated glass coverslips in a 5% CO2 incubator at 37 °C for 1 day. To investigate the effect of hIAPP on neurite outgrowth, cells were incubated with various concentrations of hIAPP (0.1-1000 nM) for 24 hours, and to investigate the effects of IAPP aggregation, 100 nM hIAPP was compared to 100 nM mIAPP, 100 nM Pramlintide, or vehicle for 24 h.

### Neurite outgrowth analysis

After incubation with hIAPP, mIAPP, Pramlintide or vehicle, DRG cells were fixed with 4% PFA for 10 mins at room temperature (RT) and washed 3 times with PBS. DRG cells were then permeabilized with PBS with 0.05% Tween (PBST) 3 times for 5 minutes at RT, incubated in blocking solution (5% normal goat serum and 1% BSA in PBST) for 30 minutes at RT, and then incubated overnight at 4°C with rabbit anti-β3-tubulin antibody (1:1500, Abcam, United Kingdom) and mouse anti-NeuN antibody (1:500, Sigma-Aldrich-MAB377). The following day, the cells were washed 3 times in PBST, incubated with Alexa Fluor 488 labeled donkey anti-rabbit antibody (1:1000, Life Technologies) and Alexa Fluor 568 labeled donkey anti-mouse (1:1000, Life Technologies) in the dark for 1 hour, washed 2x with PBST 5 min, 1x with PBS 5 min, incubated with DAPI (1:5000) 5 min at RT, washed 2x 5min with distilled water and then mounted onto slides. Confocal images were acquired with a Hamamatsu Camera C13440 Olympus IX83 microscope (Life Science) at 20x, five images randomly were taken from each well of each treatment (3 wells per condition). ImageJ with Neuralmetrics Macro plugin (46) was used to trace the neurons and their neurites automatically and measure the total length of neurite. The average neurite length/ neuron was calculated and all values were expressed as percentage of the vehicle control.

### Mitochondrial reactive oxygen species (ROS)

Cultured DRG neurons were incubated with 200nM MitoTracker deep red (Invitrogen, #M22426) and 5 µM MitoSOX Red (Invitrogen, #M36008), an indicator of mitochondrial superoxide, for 20 minutes, protected from light exposure. After washing, 4 images were taken of random areas of each well, with in total 2 wells per condition in each culture of DRG from 1 mouse. In total 3 cultures were performed from 1 female and 2 male mice. Picture were taken with an Olympus IX83 fluorescence microscope. Brightfield images were used to distinguish neurons from other cell types and select them for analysis with Image J software.

### Thioflavin T (ThT) fluorescent assay

Thioflavin T (ThT) assay was conducted in a standard 96 well black microtiter plate using a plate reader (CLARIOstar plus; BMG labtech). Per well 20 μL of each peptide (hIAPP or mIAPP or Pramlintide) was added to 180 μL buffer. The final buffer concentrations were 10 µM ThT, 100 mM NaCl, 10 mM Tris (pH 7.4). For the aggregation inhibitor experiment, per well 20 μL of hIAPP, anle145c, hIAPP + anle145c or vehicle was added to 180 μL buffer. The final buffer concentrations were 20µM ThT, 10mM tris, 150 mM NaCl, 2.5% DMSO (pH=7.4). The final IAPP and anle 145c concentrations were 12.5 µM and 250 µM respectively.

The plate was covered using a clear Viewseal sealer (Greiner) to prevent evaporation during the experiment. The plate was shaken at 500rpm for 30 sec immediately prior to the first measurement. To determine the formation of fibrils, the fluorescence intensity was followed in time. Binding of ThT to amyloid fibrils results in an increase in fluorescence (47).

The fluorescence was measured every 5 minutes (until 24 hours) at room temperature from the top of the plate with excitation at 435 nm and emission at 530nm. Measurements were performed in triplicate for each condition/concentration.

### Transmission electron microscopy (TEM)

Aliquots (4.20 µL) of 5 µM of hIAPP, mIAPP and Pramlintide were blotted on carbon coated 200 mesh copper grids, glow-discharged for 2 min. Then the samples were negatively stained with 4% uranyl acetate 2 times for 1 min. The grids were dried and examined using a FEI-CM 120 transmission electron microscope equipped with a US1900 GATAN CCD camera.

### IAPP and oligomer staining

Lumber DRG (L3-5) from WT and hIAPP mice were collected between the ages of 14 and 18 weeks, fixed in 4% paraformaldehyde (PFA), embedded in optimal cutting temperature (OCT) compound (Sakura, Zoeterwoude, the Netherlands), and frozen at –80°C.

For immunofluorescence, cryosections (10 μm) of DRG were stained with primary antibodies: mouse anti human IAPP antibody (1:500, Abcam Ab115766 mouse monoclonal), rabbit anti-I11 oligomer antibody (1:500, received from Dr. Rakez Kayed, University of Texas Medical Branch, Dept. Neurology, Galveston, USA) overnight at 4°C, followed by 2 h incubation with fluorescent secondary antibodies (Fluor 568 labeled donkey anti-mouse (1:500; Life Technologies), Fluor 488 labeled donkey anti-rabbit (1:500; Life Technologies). Images of immunostaining were captured at 20x using a Hamamatsu Camera C13440 on an Olympus IX83 microscope (Life Science).

For skin staining, samples were collected from T2DM subjects (n=6) and non-T2DM controls (n=9) undergoing surgery from hand or foot. Tissues were fixed in 4% paraformaldehyde (PFA) overnight, then embedded in optimal cutting temperature (OCT) compound (Sakura, Zoeterwoude, the Netherlands), and frozen at −80°C.

For immunofluorescence, cryosections (50 μm) of skin were stained with primary antibodies: mouse anti human IAPP antibody (1:500, Abcam Ab115766 mouse monoclonal), rabbit anti-I11 oligomer antibody (1:500, received from Dr. Rakez Kayed, University of Texas Medical Branch, Dept. Neurology, Galveston, USA) and Goat anti-collagen IV antibody (1:200, Southern Biotech, US), mouse anti Human Protein Gene Product 9.5 (PGP9.5) (1:500, MCA4750GA mouse monoclonal, Biorad) for 36 h at 4°C, followed by overnight incubation with fluorescent secondary antibodies (Fluor 568 labeled donkey anti-Mouse (1:500; Life Technologies), Fluor 488 labeled donkey anti-rabbit (1:500; Life Technologies), or Fluor 647 labelled donkey anti-Goat (1:500; Life Technologies)). Nuclei were stained with DAPI (1:5000) 5 min. Immunostaining images were captured with Hamamatsu Camera C13440 Olympus IX83 microscope (Life Science) at 10x and 40x magnification z-stack using identical exposure times for all slides. Pictures were analyzed manually for double positive (IAPP and oligomer) spots. The total length of analyzed skin was determined in order to determine the average number of IAPP-positiv oligomer spots per mm of skin. Individuals performing this analysis were blinded for patient groups.

### Statistical analysis

Data are expressed as mean ± SEM and were analysed with GraphPad Prism version 8.3 using unpaired two-tailed t tests, one-way or two-way ANOVA, or two-way repeated measures ANOVA, as appropriate, followed by post-hoc analysis. The post-hoc analyses used are indicated for each figure. A *P*-value of <0.05 was considered to indicate statistically significant differences between treatment groups/conditions. Plasma IAPP and insulin data were analysed after log transformation. The overall value of the Area under the curve (AUC) was expressed as positive by formal (-SUM (day 0 until day 4:).

### Study approvals

All animal experiments were performed in accordance with international guidelines and with previous approval from the local experimental animal committee of University Medical Center (UMC) Utrecht and the national Central Authority for Scientific Procedures on Animals (CCD) and the local experimental animal welfare body (AVD115002015323).

Human skin samples were obtained with permission from the UMC Utrecht Committee for usage of material from the UMC Utrecht Biobank, under license number TCBio-19.705.

## Data and Resource Availability

The datasets and or resources generated during and/or analyzed during the current study are included in the manuscript or in the supplementary materials and/or are available from the corresponding author upon reasonable request.

## Author Contributions

M.M.H.A., E.C.H, J.W.M.H. and N.E designed the research and performed the writing, review, and editing of the paper; M.M.H.A., B.E., J.P., and S.V. performed the experiments; E.M.B and J.H.C provided human skin tissues. M.M.H.A., J.P., and S.V. analysed the data; J.W.M.H. and N.E. supervised the work.

## Acknowledgments

This work was financially supported by King Abdulaziz City for Science and Technology (KACST), Saudi Arabia. We thank Dr. Rakez Kayed (University of Texas Medical Branch, Dept. Neurology, Galveston, USA) for providing the I11 antibody, Lucie Khemtemourian (Institut Polytechnique Bordeaux, Université de Bordeaux, Bordeaux, France) for performing the TEM imaging and Christian Giesinger, Sergey Ryazanov and Andrei Leonov (Department of NMR Based Structural Biology, Max Planck Institute for Multidisciplinary Sciences, Göttingen, Germany) for providing the anle145c.

