## Supplemental material for "Human IAPP is a contributor to painful diabetic peripheral neuropathy"

**Supplemental Data (Online Supplemental Material)**

**Table 1.** Clinical characteristics of T2DM patients and non-T2DM controls.

| **Characteristic** | **Controls**<br><br>**(n=9)** | **T2DM**<br><br>**(n=6)** |
|---|---|---|
| **Median age (range) years** | 47 (24-67) | 59 (58-72) |
| **Gender (n).** | Female (n=7)<br><br>Male (n=2) | Female (n=4)<br><br>Male (n=2) |
| **Organ** | Hand and Foot | Foot |
| **Tissue location** | Plantar and Dorsal | Plantar and Dorsal |
| **Neuropathy** | No | Yes |

Abbreviations:
T2DM= type 2 diabetes mellitus

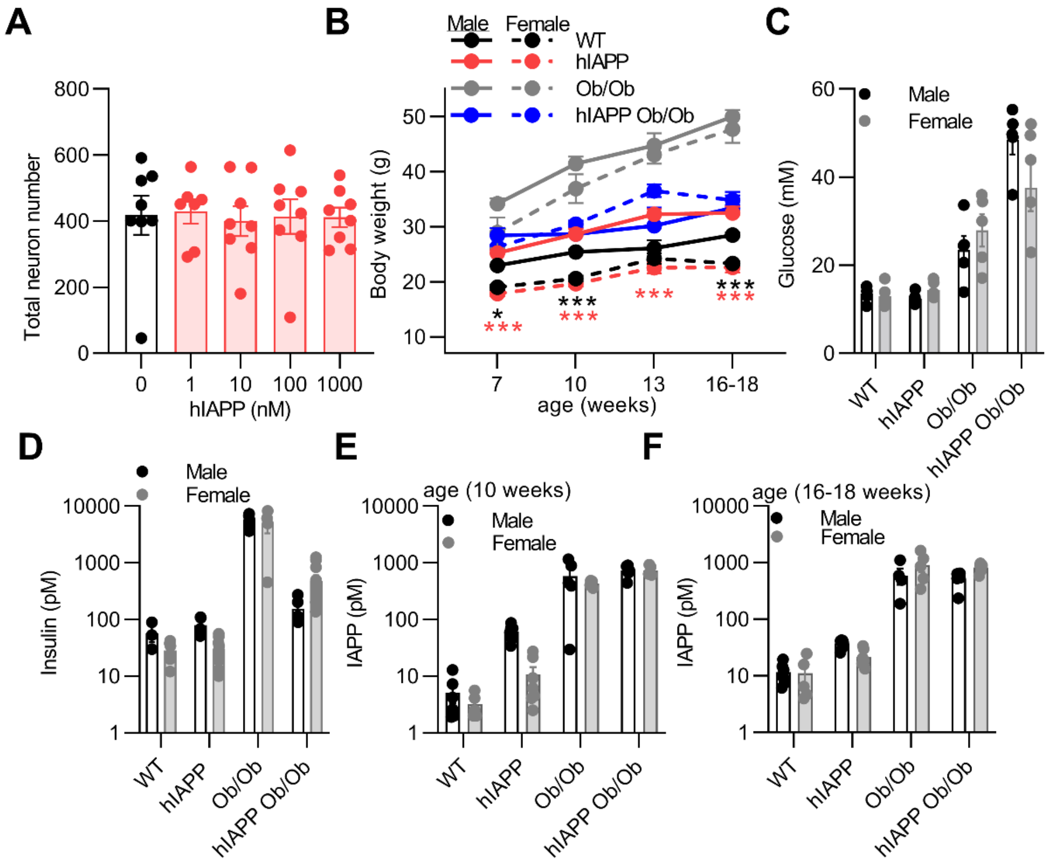


**Supplementary figure 1. Human IAPP incubation with sensory neurons and metabolic parameters in male and female WT, hIAPP, Ob/Ob and hIAPP Ob/Ob mice.** (**A**) Sensory neurons were cultured and treated for 24h with different concentrations of hIAPP (1-1000nM) or vehicle. The total neuron number was assessed (n=8; n represents a DRG culture of one mouse); **)** One-way ANOVA with Tukey's test. (**B**) Body weight of male mice (WT (n=6), hIAPP (n=15), Ob/Ob (n=16), hIAPP Ob/Ob (n=8) ) and female mice (WT (n=5), hIAPP (n=8), Ob/Ob (n=5), hIAPP Ob/Ob (n=6); Two-way ANOVA with Tukey's test; $^*p < 0.05$, $^{**}p < 0.01$,$^{***}p < 0.001$ male vs female. (**C**) Non-fasting plasma glucose level; (**D**) Non-fasting plasma insulin levels of male and female mice (**C/D;** mice age: 16-18 weeks; n=5 and, (**E, F**) Non-fasting plasma IAPP levels of male mice and female mice; n=5.(**C-F**) Two-way ANOVA with Sidak's test. Data are presented as mean ± SEM.

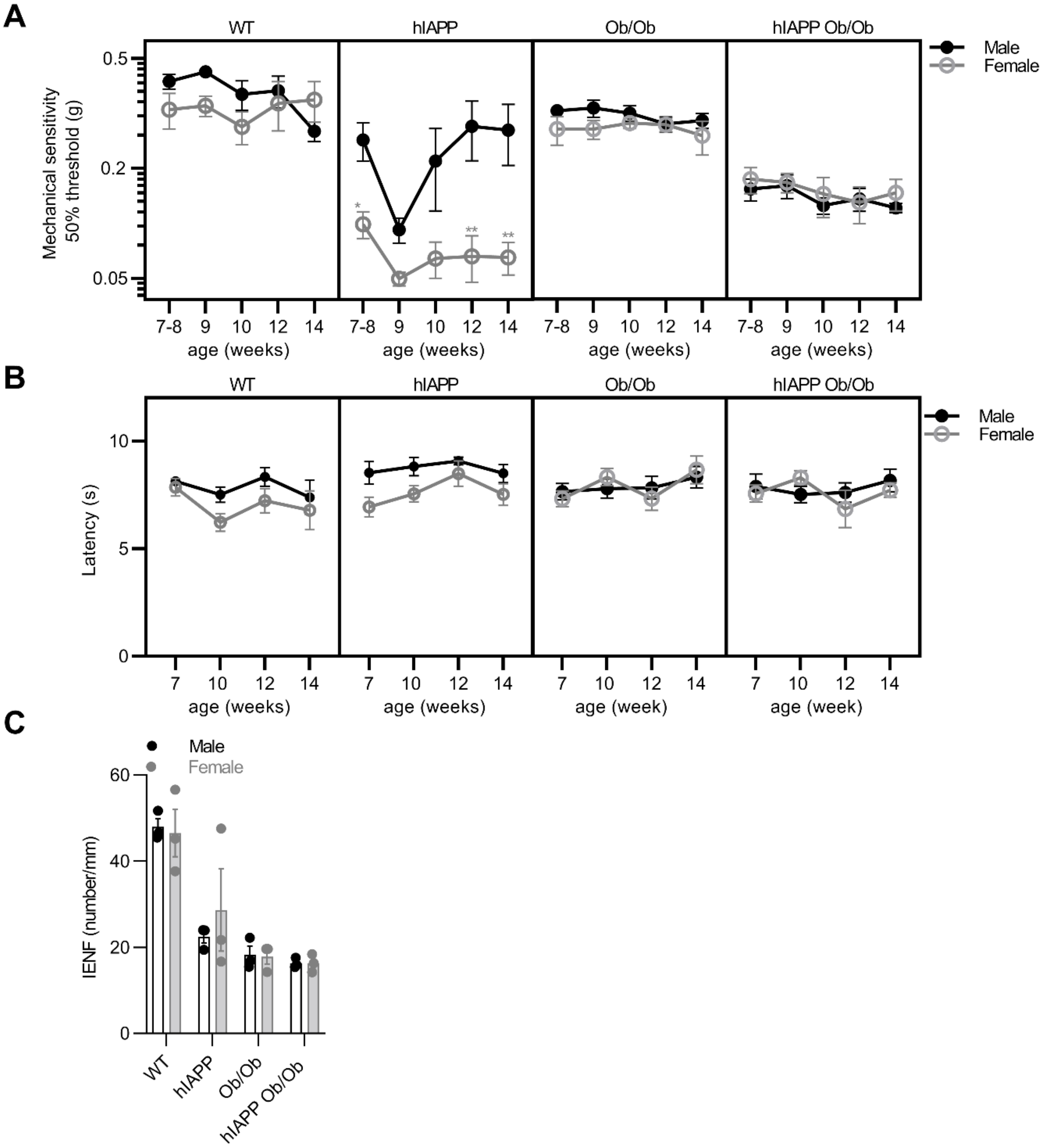


**Supplementary figure 2. Pain behaviour measurements in male and female WT, hIAPP, Ob/Ob and hIAPP Ob/Ob mice.** (**A**) Mechanical threshold of male mice (WT (n=6), hIAPP (n=15), Ob/Ob (n=16), hIAPP Ob/Ob (n=8)) and female mice (WT (n=5), hIAPP (n=8), Ob/Ob (n=5), hIAPP Ob/Ob (n=6)), Two-way ANOVA with Sidak's test; *$p < 0.05$, **$p < 0.01$ male vs female. (**B**) Thermal sensitivity of male mice (WT (n=5), hIAPP (n=7), Ob/Ob (n= 5) hIAPP Ob/Ob (n=7)) mice and female mice (WT (n=5), hIAPP (n=8), Ob/Ob (n=5), hIAPP Ob/Ob (n=5)); Two-way ANOVA with Sidak's test. (**C**) Quantification of IENF of the hind paw of male and female mice at 16-18 weeks of age; n=3 for each group; Two-way ANOVA with Sidak's test. Data are presented as mean ± SEM.

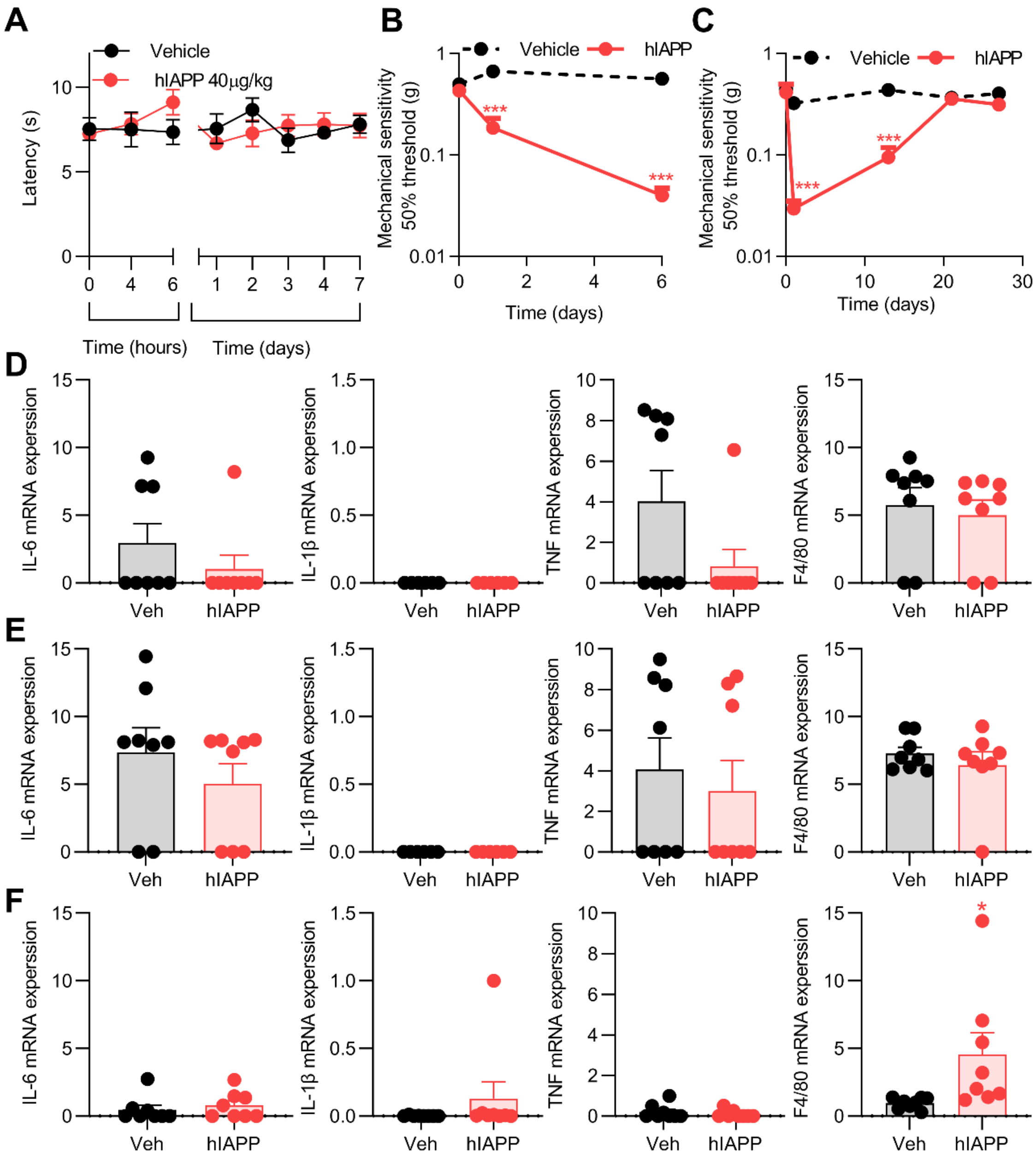


**Supplementary figure 3. Human IAPP injection in the hindpaw (intraplantar) of WT mice did not induce thermal hypersensitivity or trigger expression of inflammatory cytokines.** (**A**) Thermal sensitivity measured of the hind paw after intravenous injection of hIAPP (40 µg/kg n=6) or saline (NaCl (n=6)) into male WT mice; Two-way ANOVA followed by Sidak's multiple comparison test. (**B,C**) Mechanical sensitivity of the hind paw after intraplantar injection of hIAPP (1000fg, n=8) or saline (n=8) into male and female WT mice; Two-way ANOVA followed by Sidak's multiple comparison test; ***$P < 0.001$. (**D-F**) IL-6, IL-1β, TNF and F4/80 mRNA expression in hindpaw skin at 6 hours (**D**), 24 hours (**E**) and 6 days (**F**) after intraplantar injection of 1000 fg hIAPP or saline into WT male and female mice. Expression is normalised against housekeeping genes (average of actin, HPRT and 18S); Unpaired t test; *$p<0.05$, n=8. Data are presented as mean ± SEM.

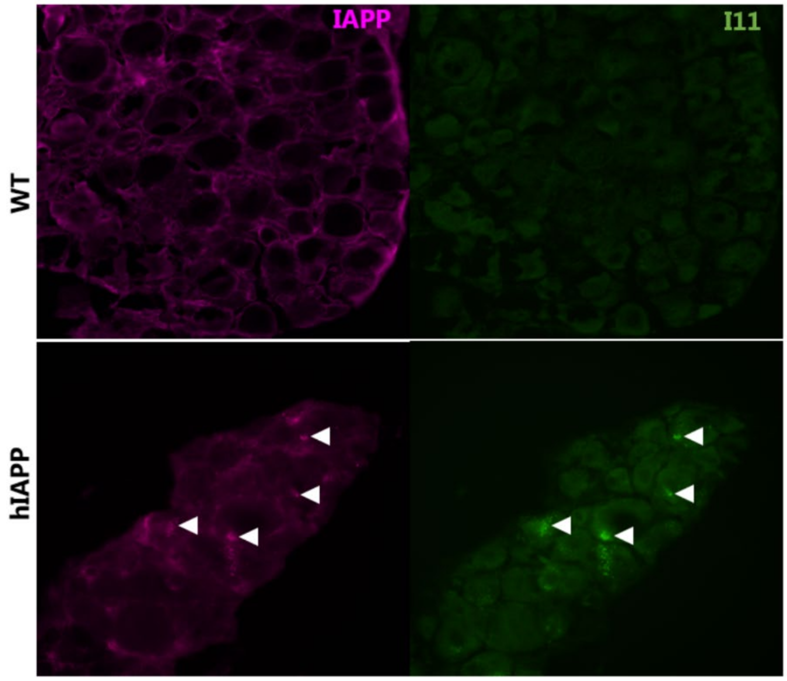


**Supplementary figure 4. hIAPP oligomers present in hIAPP transgenic mice.**
Representative images of the IAPP and oligomer staining (I11) in DRG of WT mice (controls) and hIAPP mice, IAPP and oligomer –positive spots are indicated by the arrowheads.
